# Sound-based assembly of three-dimensional cellularized and acellularized constructs

**DOI:** 10.1101/2023.05.23.541870

**Authors:** Riccardo Tognato, Romedi Parolini, Shahrbanoo Jahangir, Junxuan Ma, Sammy Florczak, R. Goeff Richards, Riccardo Levato, Mauro Alini, Tiziano Serra

## Abstract

Herein we show an accessible technique based on Faraday waves that assist the rapid assembly osteoinductive β-TCP particles as well as human osteoblast pre-assembled in spheroids. The hydrodynamic forces originating at ’seabed’ of the assembly chamber can be used to tightly aggregate inorganic and biological entities at packing densities that resemble those of native tissues. Additionally, following a layer-by-layer assembly procedure, centimeter scaled osteoinductive three-dimensional and cellularised constructs have been fabricated. We showed that the intimate connection between biological building blocks is essential in engineering living system able of localized mineral deposition. Our results demonstrate, for the first time, the possibility to obtain three-dimensional cellularised and acellularised anisotropic constructs using Faraday waves.

## 1. Introduction

Native tissue and organs present a hierarchical architecture where multiple types of cells and extracellular matrix (ECM) are intimately connected, and the structural organization is pivotal to their correct functioning.[1] Natively, biological components autonomously self-organize in highly interconnected configuration necessary for their correct functioning.[2,3] Engineering the spatial complexity of living system is vital not only to mimicry the *in vivo* architecture but also in regulating the microenvironments of the embedded cells.[4] Several biofabrication techniques have been proposed to reconfigure artificial 3D tissue surrogates.[5] In extrusion based bioprinting strategies, cells are suspended in exogenous materials and positioned by an automatic dispenser. With this technique large cellularized constructs can be obtained but with high cost in processing time and low cellular density.[6] Lately, light based bioprinting strategies emerged as valid alternative.[7] In particular, volumetric bioprinting developed as a valid method able to obtain centimeter scaled cellularized construct with unprecedent speed (few seconds) and resolution.[8,9] Nevertheless, reaching the cell density typical of the native tissue remain an open challenge.

A series of physical principles have been used to increase the local cell density such as surface energy,[10] gravity,[11] magnetism,[12] and acoustic.[13] Acoustic assembly demonstrated to be an excellent option to aggregate a large amount of cell in small area. It provides excellent biocompatibility, it is fast and contactless and enhance the spontaneous cells’ self-assembly in large functional structure.[14,15] Faraday waves are a class of standing waves emerging at the interfaces between two fluids when vibrated vertically, generally in liquid and gas phase.[16] As the vertical vibration frequency reach a critical value, the flat liquid-air interface becomes unstable, and a pattern of standing waves oscillating at half the driving frequency emerges. These standing waves generate a hydrodynamic potential field at the bottom of the liquid container. Particles lying at rest at the ’seabed’ are propelled by the potential field to migrate from region at high potential energy to region at low potential energy. This directed motion condenses particulate system at the nodal regions leaving vast region of the ’seabed’ unpopulated. Importantly, the wave patterns, and thus, the potential field, is dictated by the vibration frequency.[17,18] Interestingly, this phenomenon can be originated in liquids which can be later jellified so that the obtained particles condensate can be frozen and used on demand.[15] This sound-based assembly has been used by several researchers to investigate blood vessels formation,[19] nerve ingrowths in low back pain *in vitro* model,[20] for tumor *in vitro* model,[21] to recreate 3D *in vitro* cardiac tissue,[22] to assemble milli-scale cellu-robots,[23] and more recently to fabricate hepatic lobules.[24] Nevertheless sound-based assembly demonstrate its superior ability in creating cell condensates, the cell constructs are restricted to a 2.5D dimension because of the underlying physics. Other strategies based on holographic acoustic patterning and ultrasounds have been explored.[25–27] However, these technologies need complex apparatus and specialized personnel to operate the custom-made system, and the size of the samples are often limited to few mm^3^.

Here, we aim to demonstrate how hydrodynamic forces generated by Faraday waves can be exploited to fabricate three-dimensional osteo-inductive constructs and multilayered cellularized structures. This technology excels at producing structures with highly dense cellular composition, which is paramount to boost many cells differentiation and tissue maturation processes that are facilitated by cellular condensation and cell-to-cell contacts. To demonstrate this potential, we investigated how the localized enhancement in cells density is essential to direct a continuous and localized mineral deposition.

## 2. Methods

### 2.1 GelMA synthesis

Methacrylated groups were grafted to gelatin as previously reported.[28] Briefly, gelatin derived from porcine skin (Sigma-Aldrich) was dissolved at 10% (w/v) in PBS at 60 °C. After obtaining a clear solution, 0.14 ml of methacrylic anhydride for each gram of gelatin was added dropwise and the reaction was carried on for 3 h at 50 °C. The GelMA solution was then dialyzed against deionized water using 12–14 kDa cutoff dialysis tubes (VWR scientific) for 6 days at 50 °C to remove unreacted methacrylic anhydride and any additional by-products. GelMA was lyophilized and stored at −20 °C until further use.

### 2.2 Cell culture and spheroids formation

Primary human cancellous bone osteoblasts (hOBs) were isolated from orthopedic surgery patients according to established protocol.[29,30] In brief, bone pieces were vigorously washed in PBS and digested with collagenase type IV (Sigma-Aldrich Co.) for 45 min at 37 °C. The digested fragments were placed in 6-well plates and cultivated in DMEM low glucose supplemented with 10% FCS, 1% penicillin-streptomycin, and 50 µg/mL ascorbic acid (AA2P) and 5 mM β-glycerol phosphate (BGP).

Osteoblast micropellets were generated by seeding human osteoblasts in a 6-well microwell plate (AggreWell^TM^ 400 plate) at a density of 2×10^6^ cells/well according to the manufacturer’s protocol and cultured for one night in osteogenic medium. Cells viability within the first two layers was measured by live/dead assay with Calcein-Am/Ethidium homodimer staining. Culture medium was removed, and the cell-laden three-dimensional construct hydrogels were incubated (in 10 µM Calcein-AM and 2 µM ethidium homodimer solution in PBS for 1h at 37 °C. After the incubation hydrogel constructs were analyzed with a confocal microscope (Zeiss LSM800), equipped with a digital camera.

### 2.3 RT-PCR

Genes expressions were assessed using quantitative real-time polymerase chain reaction (qRT-PCR) at day 1, 7, and 14 of samples incubation osteogenic medium. Total RNA was extracted and purified from using Trizol reagent (Invitrogen) and RNeasy Mini Kit (Qiagen GmbH). Reverse transcription was performed with Superscript VILO cDNA synthesis Kit Manual (Invitrogen Corporation). Relative gene expression qRT-PCR reactions were set up in 10-μL reaction mixtures containing TaqMan Universal Master Mix (Thermo Fisher), the appropriate set of primers, and probes, DEPC-H2O and cDNA template. The expression of Osteopontin, Osteocalcin, SP7, Coll I, ALP, and Runx-2 were evaluated and PRLPO was used as a housekeeping gene. Primer and probe sequences are shown in supplementary data Table S (supplementary material), whiles catalog numbers of Assays-on-Demand (Applied Biosystems) are listed in the supplementary data Table S (supplementary material).

### 2.4 Acellularized and cellularized construct imaging

Cell laden three-dimensional construct was imaged using a custom-built light sheet microscope at a 532nm excitation. The computed tomography scans were obtained using the cone-beam microCT scanner VivaCT40 (SCANCO Medical AG). A 21.5 mm field of view, at a voltage of 70 kV and 114 µA current, 200 ms integration time, and 1000 projections per scan were chosen. The projections were then reconstructed across an image matrix of 2048 × 2048 pixels resulting in an isotropic voxel size of 10.5 µm.

### 2.5 Data and image analysis

Image analyses were performed with ImageJ Fiji (1.53q NIH). Data analysis and simulation were performed using custom made script (Python Version 3.9 and Numpy Library).

## 3. Results and Discussion

Experiments can be conducted with a simple and cost-effective setup. Here, we used an experimental configuration like the one previously reported by Petta *et al.*[19] A sample holder (framed in aluminum) accommodates a commercially available petri dish. Within the petri dish a custom-made plastic frame (PMMA) containing the liquid receptacle/receptacles can be lodged. The sample holder is then connected to an electromagnetic mechanical driver (Pasco Scientific) that transfers sine wave signals to the liquid by a vertical vibration at specific frequencies (Fig. 1-A). Generally, the assembly procedure can be resumed in few steps. A particulate system is mixed with a liquid solution of a pre-polymer and then is loaded in the liquid receptacle (Fig. 1-B) that is later vibrated vertically by the mechanical driver (Fig. 1-C (i)). Once the particles are assembled in the desired pattern, the polymerization of the pre-polymer solution is triggered by a specific stimulus (e.g., temperature, UV-Light, enzyme, etc.), the assembled structure is frozen in the acquired shape (Fig. 1-C (ii)), and therefore, the sample can be retrieved for further use, (Fig. 1-C (iii)).

**Figure 1.**
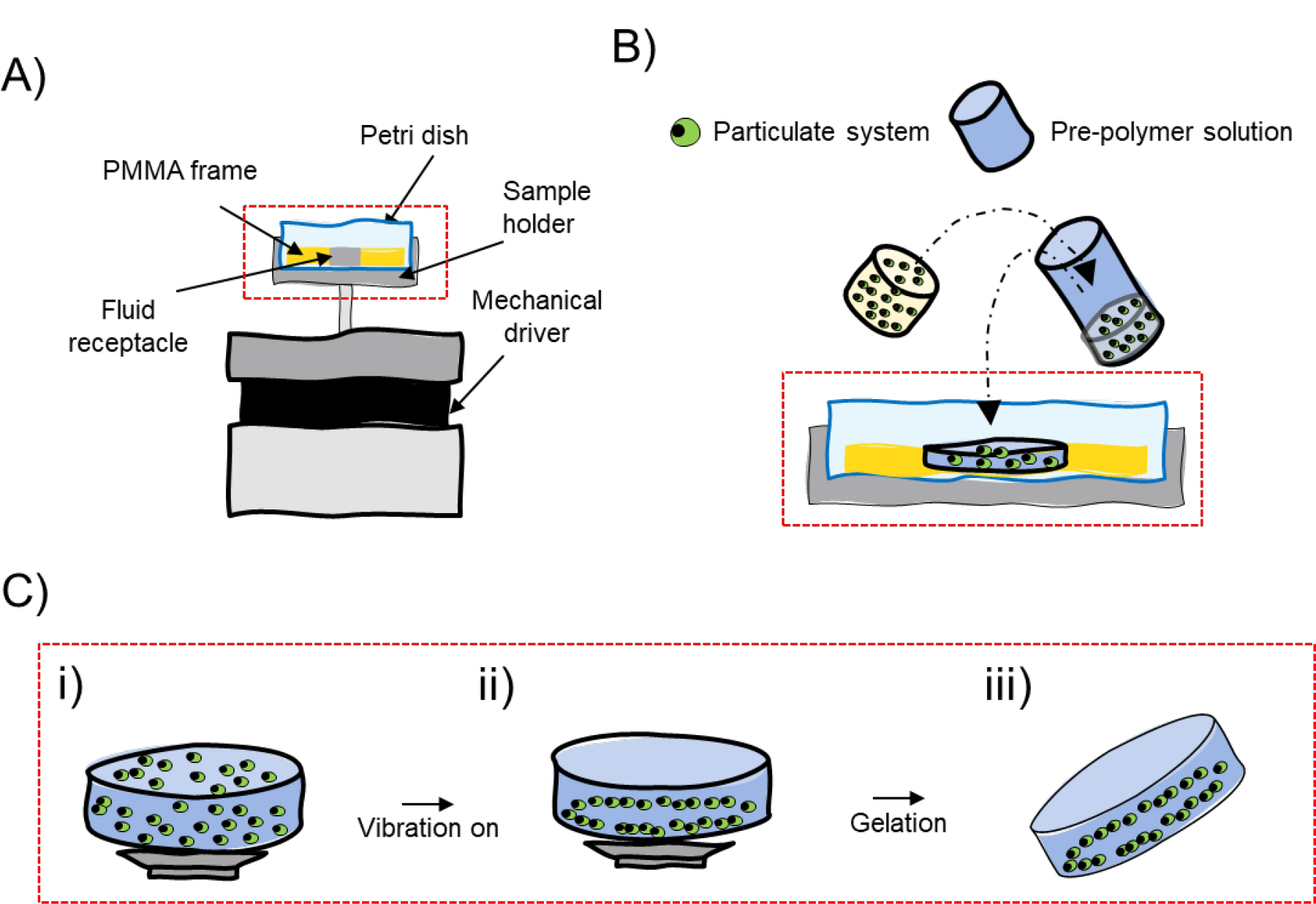
Experimental setup and assembly procedure. Schematic depiction of the experimental setup (A), sample loading (B) and acoustic assembly steps (C): i) the liquid GelMA-particles dispersion is disposed in a liquid receptacle. ii) The liquid dispersion is vibrated vertically at specific frequency exciting specific Faraday waves pattern which guides the assembly of the particulate system in specific nodal zone. iii) The prepolymer solution is cross-linked freezing in place particles’ pattern which can be retrieved for further use.

Initially, we tested our setup by assembling β-TCP microparticles (35-75 µm in diameter, Kuros bioscience) in the stimuli responsive gelatin methacryaloyl (GelMA 5% w/v, see material and methods for the detail about the grafting procedure). GelMA presents a phase transition temperature of ≈30 °C (being liquid above and solid below), good biocompatibility, and can form an irreversible, covalent gel via the free-radical polymerization of the acryloyl groups, which can be obtained, for instance through crosslinking via visible or UV-light in presence of a photoinitiator (*i.e.*, as used in this study, Lithium-Phenyl-2,4,6-trimethylbenzoylphosphinat, LAP, Sigma-Adrich).[28] β-TCP microparticles particles were dispersed in pre-warmed GelMA (5% w/v in PBS and 0.1 % LAP) and assembled in the simplest possible shape: a toroid.

Toroidal structures offer a series of advantages in characterizing the experimental setup and the fidelity of the assembling procedure. A ring shaped (or multiple concentric rings) structure can be analyzed through the image processing proposed by Di Marzio *et al.*, performed with ImageJ Fiji (1.53q NIH, US) and custom-made scripts (Python 3.9 and numpy/scipy library)[21]. After imaging the obtained structure, the intensity of the pixel along every circumference of a defined radius can be summed and plotted against the specific radius (radial profile plugin authored by Paul Baggethun). For ring-like structures, this analysis originates a plot with a precise peak positioned at a specific distance from the center of the image, provided enough contrast between the particles and the background.[21] Radial profile analyses of the images of multiple assembled constructs shows a single peak sharpening at ≈4 mm from the centre of the structure indicating the accumulation of β-TCP in a ring Fig. 2-A and -B. Numerical simulation of the surface displacements (Fig. 2-C) and hydrodynamic drag forces (Fig. S1-B(i)) show that β-TCP accumulate in location where the force potential is lowest and that are distributed as a ring with a diameter of ≈4 mm Fig. 2-D.[23] Importantly, this analysis is not limited to single images obtained at the final time points but can be fully automated to analyze time lapse video of the assembling process obtaining useful parameters regarding the assembly kinetic . Fig. 2-E shows the evolution of the assembly process when analyzed with the radial profile. Initially, when the dispersion is at rest, no peaks are evident but within 2 s a toroidal structure appeared, Fig. 2-A. This can be quantitatively measured by calculating the area under the peak obtained at each time point. In Fig. 2-G are reported the values of the areas under the peak as a function of the assembling time. Here, it is visible, that after the appearance of the surface waves the assembled structure forms within 1 s (Fig. 2-F), given specific particles and fluid properties. These data are well fitted with a logistic function since the area under the peak reaches a plateau when the structure is completely formed. Averaging over multiple repetitions we obtain an assembling rate of k = 7.102 ± 2.452 1/s (Fig. S2). This procedure is general and can be extended to different particulate material and suspending medium/prepolymer solution. Differences in the assembly rate can be observed when fluids of different viscosity are used as dispersing medium (Fig. 2-F). Generally, in less viscous fluid the assembling is faster as observed by the steepness of the logistic growth (Fig. 2-F).

**Figure 2.**
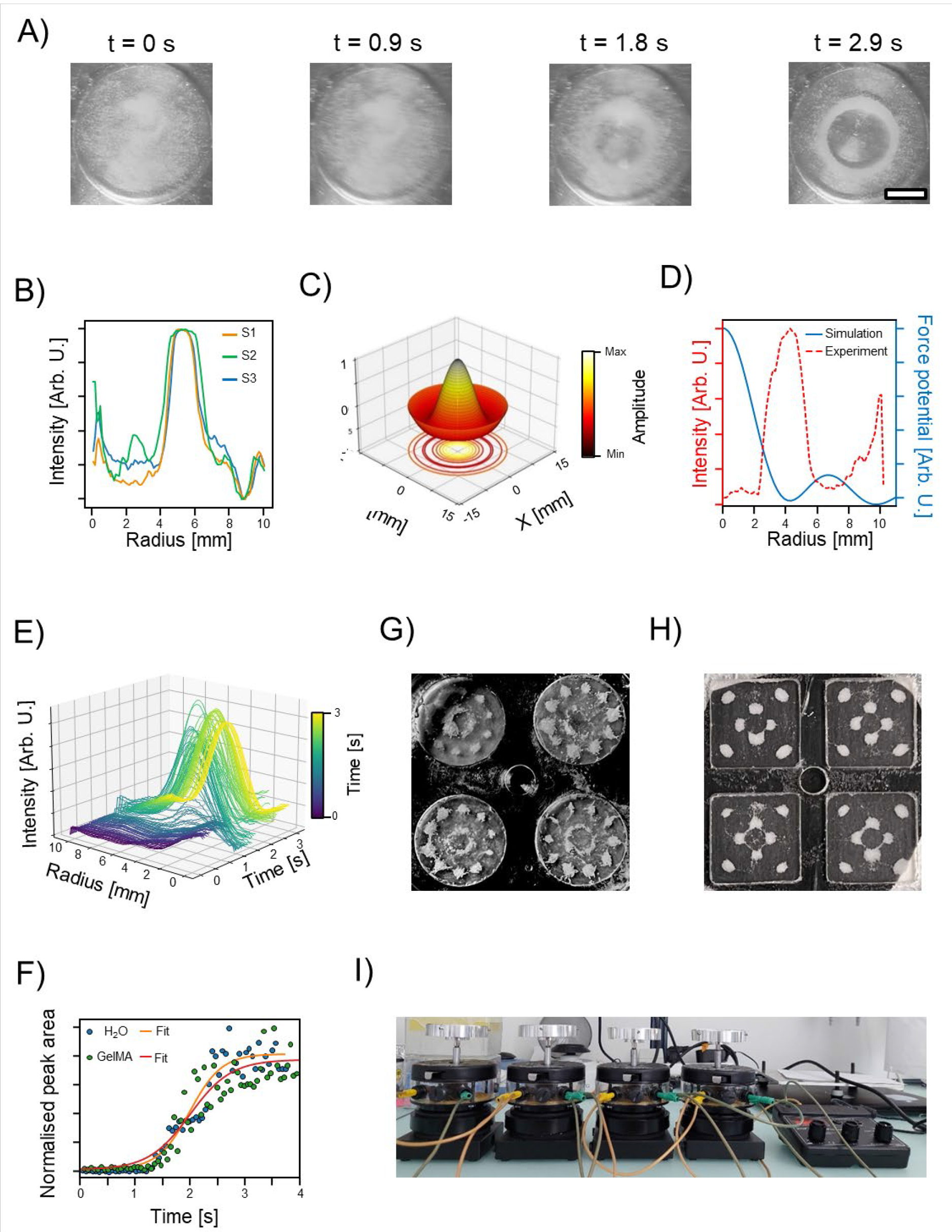
Optimization of the assembly process for 2D osteoinductive constructs. A) Time- lapse images of the acoustic driven condensation of β-TCP particles in GelMA 5% w/v. Scale bar = 5 mm. B) Radial profile analysis of multiple samples compared for reproducibility. C) Numerical simulation of the fluids interface deformation driven by Faraday wave. D) Quantitative comparison between experiments and simulation for an assembled single ring. E) Radial profile analysis over time and kinetic of the assembly process in (F). Parallel assembly process in round (G) and squared chamber (H), and assembly procedure using multiple mechanical drivers (I).

Successively, we tested the possibility to assemble multiple samples at once with a single mechanical driver. We designed a single frame with multiple (four) liquid chambers with either squared (14 mm side, 1.5 mm thick) or round (13 mm diameter, 1.5 mm thick) boundary (Fig. 2-G and Fig. 2-H). In both cases, we were able to assemble the same structure in multiple chambers with a higher level of complexity compared to the simple toroid shown for the single chamber (≈100 Hz for circles and ≈69 Hz for square chamber). It should be noted the frequency of vibration at which the abovementioned structures could be formed, is specific for the type of chamber and fluid used in the experiment. The optimal frequency, in fact, varies as a function of key experimental parameters, including chamber size, fluid viscosity, particle type, sample holder, etc. Also, the liquid receptacle must be filled until the nominal volume (or slightly overfilled) to avoid the formation of concave meniscus at the boundaries that would give rise to capillary forces that pull the particles towards the edges. Later, we assessed the scalability of our process. We evaluated the possibility of assembling multiple samples using four mechanical drivers that excite a variable number of liquid chambers each controlled by a single wave generator (Fig. 2-I). With this system, we were able to assemble β-TCP in geometry ranging from simple toroid to more complex shape. The number of samples that can be prepared at once is only limited by the design of the sample holder and the liquid receptacles. In fact, this parallel assembly procedure can be further scaled up to hundreds (probably even thousands) of samples by simply designing sample holder that can accommodate tens of liquid receptacles each or connecting in parallel more machine.

Sound-based assembly through Faraday waves presents several advantages (e.g., fast, contactless, mild, etc.) but, because of the underlying physics, it typically allows to fabricate only 2.5-D constructs.[17] Having a solid ’seabed’ is essential for this assembly strategy. To excite the specific wave-mode, the vibration generated by the mechanical driver must be transmitted to the superficial fluid by the basal layer. Moreover, a particulate system must lie at the bottom of the assembly chamber to be assembled by the hydrodynamic forces in a stable position. This led us to consider only two possible working options to obtain 3D constructs: preassembling different layers and embedding them in a second hydrogel or proceeding with a layer-by-layer assembly strategy. We decide to overcome this limitation by proceeding with a sequential procedure as schematically depicted in Fig. 3-A. In complete analogy with the 2D case, one starts by mixing the particulate system with a prepolymer mixture adding it to the fluid receptacle, Fig. 3-A (i). Therefore, a first structure is assembled in the dedicated insert glued to the bottom of a commercially available petri-dish. Once the structure is assembled, a partial cross-linking for 2 minutes to solidify GelMA through UV-light is perfomed (Bio-Link BLX-365UV radiation chamber, 0.1 J cm^−2^), Fig. 3-A (ii). Therefore, a second insert is manually aligned and glued on top of the first one, and the β-TCP GelMA suspension is added to the chamber and the structure assembled, Fig 3-A (iii). Once the second layer is formed a third can be assembled on top of the previous one. This process can be repeated until the desired number of layers is reached. The final construct is then completely cross-linked for 10 mins to connect the different layers and retrieved for further use, Fig 3-A-(iv). While the process as we performed in this experiment is labor intensive, and the precision of the operator could limit the number of accurately sequential layers obtainable, this practical limitation could be overcome, for example by pairing the device with robotic arm, or more generally with an automated placing system.

**Figure 3.**
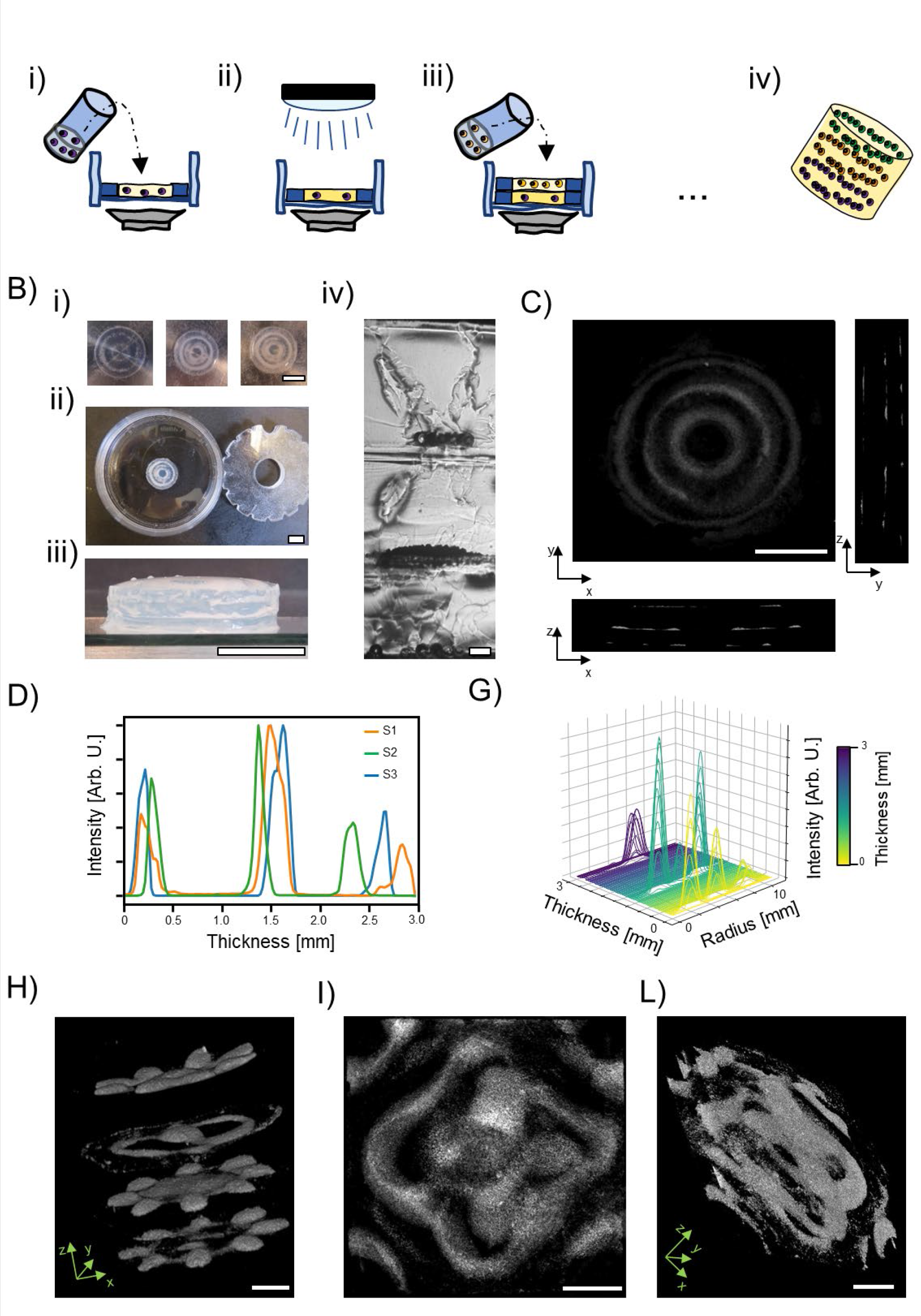
Fabrication of 3D osteoinductive constructs. A) Schematic depiction of the 3D construct fabrication procedure. The structure is fabricated by a layer-by-layer assembly and partial cross-linking assembly procedure (i-iii) of several GelMA layers. As the final thickness and morphology is reached the partial cross-linked structure is completed with UV-light and the structure retrieved for further use (iv). B) Pictures of the layer-by-layer fabrication procedure (i) top view (ii) and side view (iii) and cross-section section of the final construct (iv). Scale bars = 7.5 mm (i-iii) and 200 µm in (iv). C) Computed tomography (CT) reconstruction of ’Christmas tree-like’ structure. Scale bar = 6 mm. D) Normalized pixels intensity profile of slices along the thickness of the constructs shown in (C), and radial profile of pixel intensity of each single plane obtained from the CT imaging. CT reconstruction of 3D constructs assembled in circular (H) and squared (I-L) chamber. Scale bar = 3 mm.

Next, we verified our procedure using two particulate systems with different physical properties (β-TCP, and plain polystyrene particles 100 µm diameter). Also, to increase the complexity of our system, we choose to pattern the particles in two concentric toroids in three overlapping successive layers. To achieve this, we use a frequency of ≈66 Hz for every layer. We observed that the amplitude of the vibration must be tuned for every layer, generally increased as the thickness of the construct increases (e.g., solution viscosity, etc.). Fig. 3-B shows pictures of the entire procedure. The bottom layer is composed by two concentric toroids of plain polystyrene particles (Fig. 3-B (i) left). After that, we assembled a second layer of inorganic β-TCP and subsequently a third layer of plain polystyrene microspheres, Fig. 3-B (i). The radial profile analysis confirms that for each layer two sharp peaks are present at ≈3.2 and ≈7.5 mm, values in agreement with the numerical simulation (Fig. S4 and Figure S1 A-C (ii) respectively). The complete overlapping of the patterned particles is also confirmed by a cross-sectional image where the nature and the position of the particle is easily imaged, Fig. 3-B (iv). Not surprisingly, when completely cross-linked the entire construct can be easily removed from the assembling chamber and further used, Fig. 3-B (ii-iii).

Now, we show that the same type of particles can be patterned with a different structure in 3 successive layers. We kept our model of concentric toroids and β-TCP particles, opting for a ’Christmas tree-like’ structure where the number of toroids decrease from three to one moving from the lowest to highest layer. To drive the assembly in the specific shape we used frequency of ≈110, ≈66, and ≈28 Hz for the first, second and third layer respectively. Fig. 3-C shows the maximum projection and cross-sections of a computed tomography (CT) of the entire construct retrieved from the inserts. In Fig. S5-A the entire structure is 3D reconstructed. From these, it is clearly visible a particulate system with increased scattering properties and well-defined structures compared to that of the surrounding materials. Importantly, it is clearly visible that the particles are assembled in concentric toroids in every layer, Fig. 3-C. To extract quantitative data, we analyse the total pixels intensity (and its spatial distribution) in each CT slice. Fig. 3-D shows the normalised integrated pixels intensity of each slice of the CT scan, for three experimental repetitions (S1, S2, S3, respectively). For each sample, three peaks are clearly visible representing the plane where the β-TCP are assembled. The peak’s maximum can be assumed where most particle are assembled in the z-direction and can be used to calculate the interlayer distance. The peak-to-peak distance is 1.015 ± 0.063 mm, value is in perfect alignment with the thickness of the receptacles (1 mm). However, for the last layer (peak at ≈2.5 mm), the profiles showed lower degree of alignment, when compared to the first and second layer. This could be completely related to the operator errors in aligning the inserts during the fabrication procedure. Later, we also evaluate the pixels’ intensity distribution in each slice over the entire thickness of the sample. As previously said, toroidal structures are particularly well suited to be analysed with the radial profile procedure. Here, we extended the procedure shown in Fig. 2-E to the third spatial dimension (thickness of the construct) instead of time. Fig. 3-G shows the three-dimensional radial profile for a construct with ’Christmas tree-like’ conformation. Starting from the basal layer (thickness = 0 mm) we see 3 clear peaks, 2 of similar intensity in the second, and 1 in the last layer. We attribute the third peak in the second layer to some debris that stuck irreversibly to the layer beneath. Nevertheless useful for optimisation process, one could think that our technology is limited to the possibility to obtains toroids with at most 3 layers in circular receptacle. Therefore, we decided to highlight the possibility to pattern different structures over 4 layers (Fig. 2-H) where three layers show the same pattern, ’flower-like’ structure, while one shows a different structure ’Tao-like’ structure. Also, in Fig. 3-I and L, we show the 3D CT reconstruction that demonstrate our ability to pattern structure with defined shape in 3D utilising squared chamber. Even if not shown in the paper, we would like to highlight that a similar procedure and three-dimensional patterning could be possible also with fluid receptacle with shapes that deviates from square and circle.

Established a protocol for the precise fabrication of three-dimensional osteo-inductive scaffold, we focused our attention on the fabrication of cellularized constructs. We hydrodynamically assembled spherical multicellular aggregates of osteoblast (mean diameter 125 µm, Fig. 3-A), using GelMA as polymeric matrix. GelMA demonstrated to be a good supportive material for osteogenic differentiation and calcium deposition already at concentration 5% w/v.[31] Thus, cells’ spheroids (3000 spheroids/ml) were assembled in donut-like structures with approximately ≈3.5-4 mm radius as highlighted by the micrograph and the radial profile Fig. 4-B (i) and -C respectively. These values agree well with the numerical simulation, Fig. 2-D. Cells growth was monitored over 7 days of culture through image analysis of the tile reconstruction. Radial profile analysis can be used to monitor the radial sprouting of cells. In fact, as the radius of the circular annulus increases, the full width at half maximum (FWHM) of the radial profile plot increases as well, as shown for the simulated data in Fig. S4. Precisely, radial profile was measured at day 1, 3 and 7, observing a linear increase of the FWHM. This is also accompanied by a linear increase of the total area covered by cells over time, Fig 4-D. Together, these results suggest a radial sprouting of cells from the initial assembled ring towards free regions where cells/spheroids were not densely packed. Such observations are in accordance with previously reported results.[19,22] In addition, this continuous and pronounced cells’ sprouting highlights once more the mildness of the assembly procedure.

**Figure 4.**
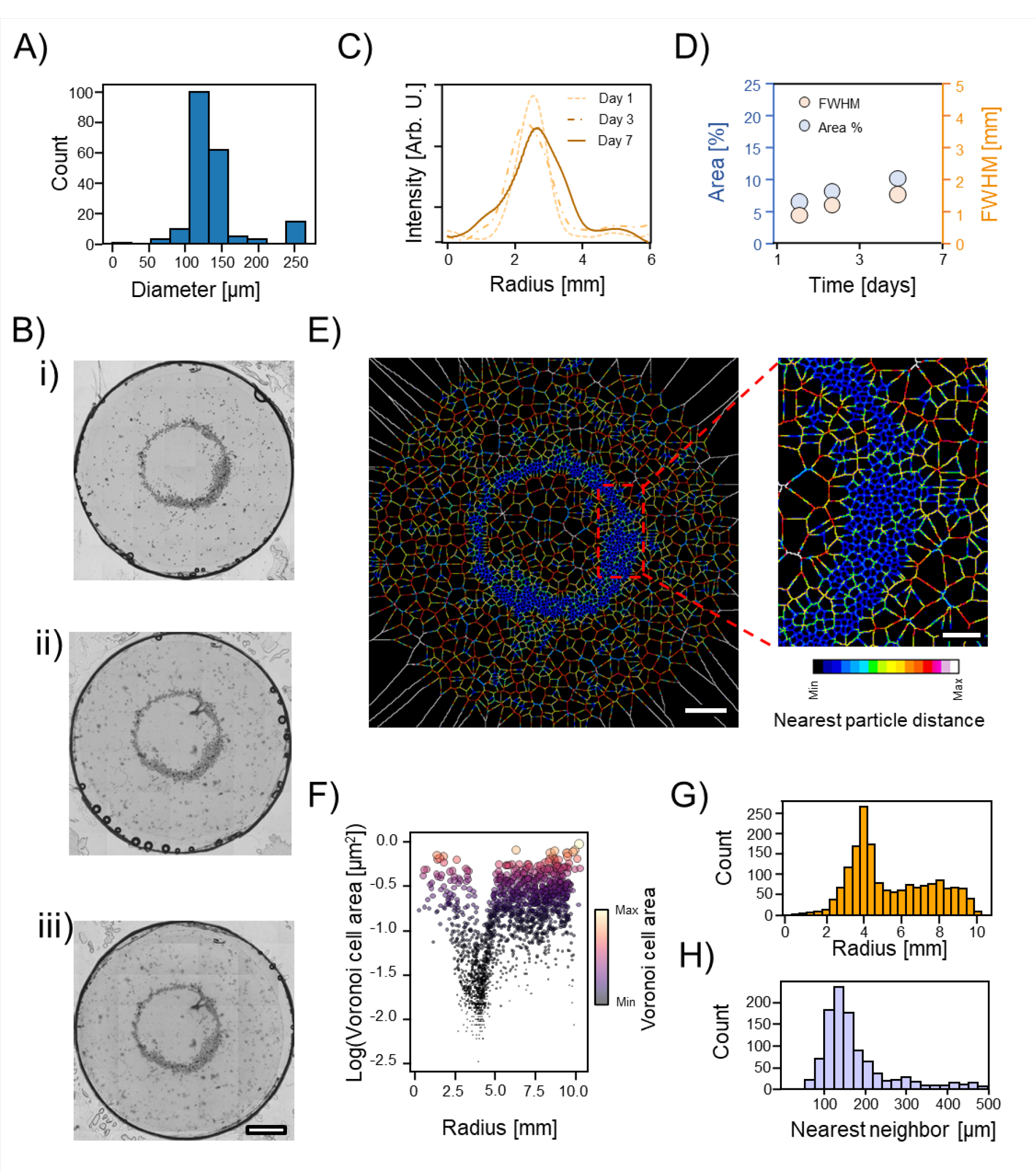
Acoustic assembly of osteoblast spheroids. A) Size distribution of osteoblast spheroids. B) Cellular aggregates assembled in toroidal structures after one (i), three (ii) and seven days (iii) of culture. Scale bar = 2 mm. C) Radial profile estimation for the osteoblast after 1, 3, and 7 days of culture and quantification of the cell spreading area and full width at half maximum (FWHM) of the radial profile traces (D). The darkest pixels were use as seeds to calculate the Voronoï tessellation map (E, scale bar = 2 mm and 0.5 mm respectively), estimate the available area to each spheroid as function of the radial distance from the center of the toroid (F), spatially count the cellular aggregate (G) and calculate the nearest neighbor to each single biological particle (H).

Later, we quantified the packing degree of the cellular spheroids within the toroidal structure via image analysis. Starting from the images shown in Fig. 4-B, we invert the pixels values, and we identify the brightest pixels which correspond to the darkest spots in the optical images, Fig. 4-B. These spots are used to locate the spheroids center in 2D for the successive analysis. Therefore, these seeds are used to build the Voronoi map of the 2D space, Fig. 4-E. The edge of the Voronois’ tessels are color coded to reflect the distances between two seeds. In Fig. 4-G, it can be seen that most spheroidal particles are packed in an anulus with radius ≈3.5-5.5 mm from the center of the toroid. Within this anulus (Area = 56.549 mm^2^), the area available to each spheroid decreases sharply compared to the spheroids’ area outside the toroidal structure, Fig. 4-F. In fact, when seeds were used to calculate the nearest neighbor of each particle, a sharp peak appear at ≈120-130 µm which is approximately the same the diameter of a single spheroid demonstrating that most of the particulate system is in tight contact between each other over a vast area. This shows the possibility to precisely manipulate organize biological entities across scales.

We demonstrated the ability to assemble biological entities over a vast area mimicking the tight contact present in the native tissue. Next, we exploited this mild technology to engineer a living system for the localized mineral deposition. We decided to adopt a more complex pattern and to shift to a squared liquid receptacle, while we kept our model cells and polymeric matrix. Single cells were initially assembled in spheroids overnight (see methods sections) obtaining spherical cellular aggregates of around ≈125 µm in diameter. Later, the spherical particles were recovered and dispersed in pre-warmed GelMA solution (5 % w/v, 0.01% LAP). Therefore, the suspension is added to the squared receptacle and the vertical vibration applied (≈98 Hz). Within few second spheroids assembled at the nodal region creating the cross-circle pattern shown in Fig. 5-A. The assembled structure was stabilized via UV cross-linking for 10 mins. Despite we created a more complex pattern, cells grow well and extend outside the initial pattern over 7 days of culture in osteogenic medium as demonstrated by the linear increase of the area covered by the cells, Fig. 5-A and -B. After, we evaluated the expression of early osteogenic markers between a patterned and a non-patterned control over 14 days of culture. We did not observe any difference between the assembled samples and the random control in terms of genes expression over time, as shown by the RT-PCR in Fig. 5-C. However, we do believe that our ability to locally enhance the cell density to a direct cell-to-cell contact could pivot the localized mineral deposition at the centimeter scale. Yet, in a random control, the same cell density and direct cell-to-cell contact can be reached increasing the number of spheroids to completely cover the receptacle area, this would make us lose the ability to obtain a localized and anisotropic cellularised structures. For this reason, after we verified that no major biological response is triggered by the enhanced cell density, we exclude the random control from the next experiments. With this in mind, we evaluated the calcium deposition of our construct after 14 days of culture in osteogenic medium. In Fig. 5-D is shown the reconstruction of the structure stained with alizarin red following a previously reported protocol.[32] Here, it can be seen that the biggest calcium deposits were colocalized with the assembled osteoblast spheroids, as confirmed by the comparison of the radial profile measurement carried out on the 4 small circles, Fig. 5-E. From a closer inspection, it is clear that the biological activity (calcium deposition ability) of each spheroid is unchanged once patterned, but the close proximity between the cell aggregates (∼150 µm) makes the inter- spheroids communication easier. Calcium can be locally deposited where the osteoblast spheroids were locally aggregated, Fig. 5-D close-ups. By contrast, this local and anisotropic mineral deposition would be impossible when spheroids are randomly dispersed in the polymeric matrix at the same density since the average distance between the single spheroids would much higher. Moreover, as previously suggested by Ren *et al.*, the aggregation of cells in thin structures (*e.g.*, toroids, lines, etc…) increases the surface area to volume ratio (*i.e.*, shrinkage of the cellular constructs along one or more spatial directions) while preserving the cell condensation and thus the biological activity.[33] Interestingly, this enhances the influx/efflux of nutrients and metabolic byproducts from the inner core of the cellularized assembly decreasing, at least to some extent, the need of vasculatures.

**Figure 5.**
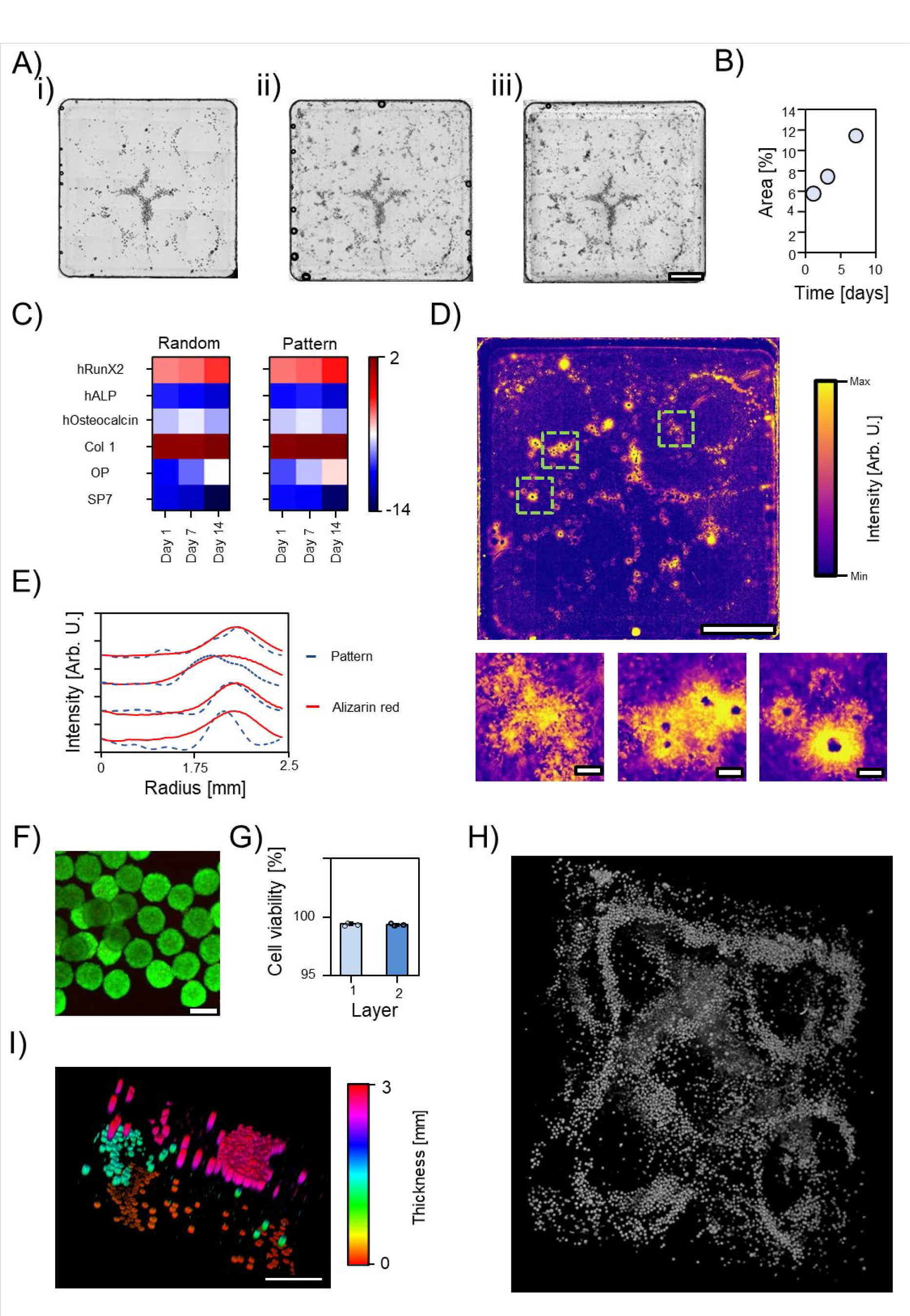
Localized mineral deposition and three-dimensional patterning. A) Cellular aggregates assembled in cross-circles structure after one (i), three (ii) and seven days (iii) of culture. Scale bar = 2 mm. B) Quantification of spreading area. C) Heat map of RT-PCR data comparing the patterned and the random sample after 14 days of culture shown as log2 of fold change. D) Alizarin red staining of the patterned cross-circle structure after 14 days of culture in osteogenic differentiation medium. Scale bar = 2 mm and 200 µm. E) Radial profile comparison between alizarin stained and unstained sample shown in (A). F) Live-dead staining of the basal layer of a multilayered sample at day 0 and (G) viability quantification of the two basal layers. H) Three-dimensional reconstruction of a cellularized multilayered sample with a total volume of 1.2×10^3^ mm^3^ and a confocal imaging of a section of the entire construct (I, scale bar = 1 mm).

Hydrodynamic forces can be exploited to obtain three-dimensional constructs by a layer-by-layer patterning. We followed the same approach used for the inorganic particles to obtain a three-dimensional living system. Human mesenchymal stem cells (hMSCs) were cultured as previously described.[34] hMSCs spheroids were prepared as described in the methods section, embedded in GelMA (5% w/v and 0.01% LAP) and disposed in the assembly chamber. The vertical vibration was applied until cellular aggregates assembled at the nodal region. The structure was therefore partially cross-linked with UV-light for 2 minutes. The second layer was patterned on top of the first one. Once the second layer was assembled, a third layer was fabricated on top, and the structure was completely cross-linked for 10 mins. This approach was successfully used to pattern spheroids on multiple layers with squared and circular chamber, Fig. 5-I and -H. After one hour of culture, cells viability was evaluated via live and dead staining. Cells’ viability within the layers most exposed to UV-light (first and second) was not compromised as shown Fig. 5-F and -G. Crucially, in this three-dimensional architecture, the cell-to-cell (or spheroids-to-spheroids) communication is maintained within each single layer. Cells could migrate between different layers establishing new connections, and creating a unique tissue, eventually. Also, this modular architecture could be used to recapitulate tissues interfaces to study cells invasion *in vitro*. Moreover, the matrix chemical composition and mechanical properties could be exchanged allowing a better modulability to simulate tissues on demand. However, it must be stressed that currently the different layers are still separated by big distance (1 mm), and this is dictated by the chamber thickness. In future, other methodologies such as magnetic levitation,[35] could be coupled with the sound-based assembly to obtain more relevant construct.

## Conclusion

In summary, we showed a methodology to fabricate anisotropic osteoinductive constructs patterning β-TCP microparticles in predetermined shape in 2D in few seconds with high reproducibility and high fidelity. The technique allows for an easy and rapid scale-up and the simultaneous fabrication of multiple scaffolds by designing inserts accommodating multiple receptacles and/or parallelizing multiple mechanical drivers. Further, we demonstrate for the first time that the sound-based assembly based on Faraday waves can be used to create osteoinductive three-dimensional construct with β-TCP finely organized with different architectures within up to four superimposed layers. Also, we proof the efficacy of the sound-based assembly in condense cellular aggregates over an area at centimeter scale, where the intimate spheroids-o-spheroids contact is maintained. The maintained close interconnection between biological entities over a vast area paved the waves to engineer a living system able to locally deposit minerals.

## Acknowledgements

Project no AOCMF-21-04S was supported by AO Foundation, AO CMF. AO CMF is a clinical division of the AO Foundation - an independent medically-guided not-for-profit organization.

## Supplementary information

**Figure S1.**
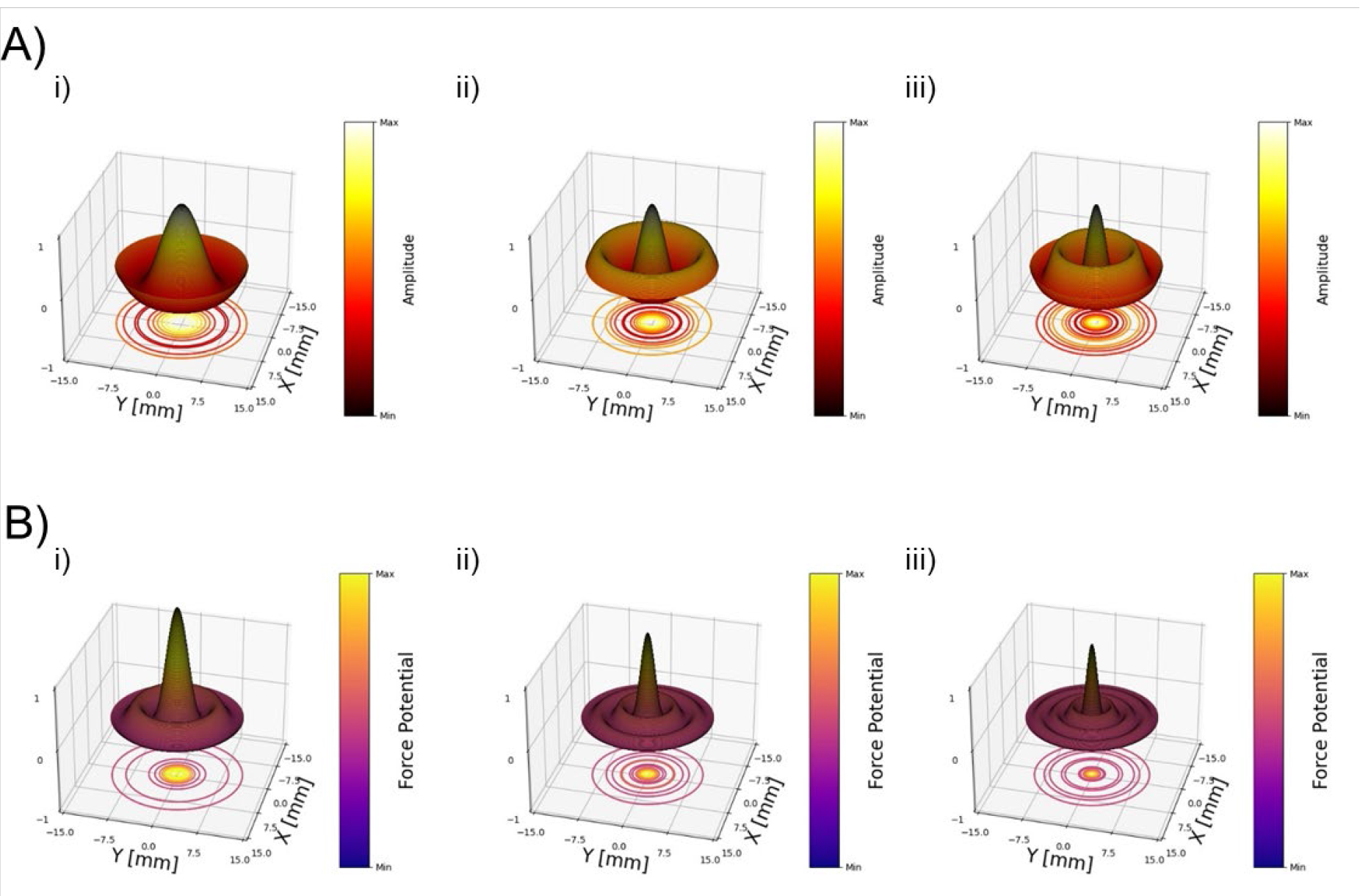
Numerical simulation surface displacements (A) and force potential (B) for one (i), two (ii) or three (iii) nodal circles. Simulations are based on the work proposed by Lei *et al.*and Ren *et al.*[2]

**Figure S2.**
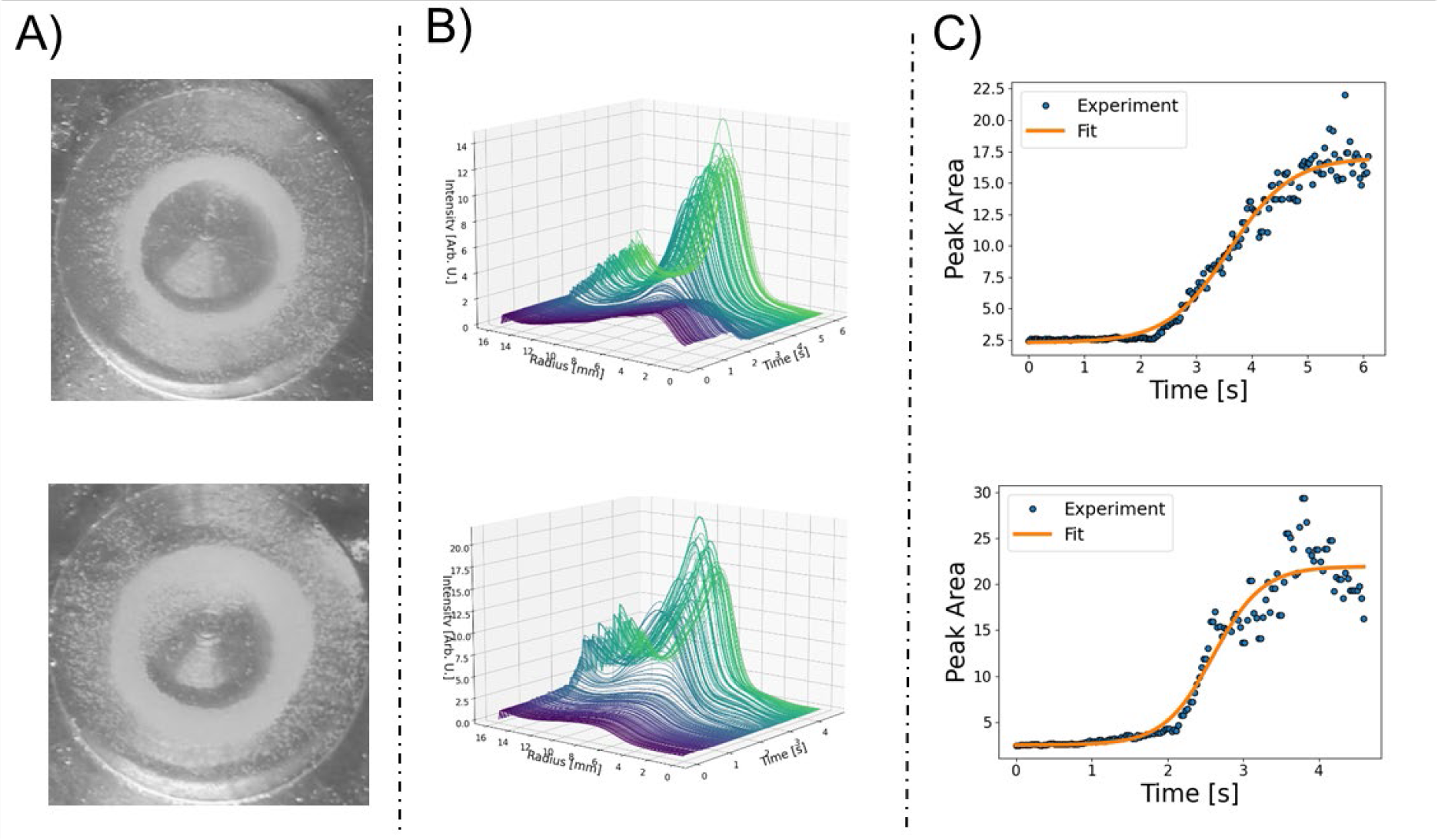
Repetition of the assembly kinetic of β-TCP particles in GelMA 5 % w/v. (A) Final structure, (B) radial profile analysis over time and (C) curve area as function of assembly time.

**Figure S3.**
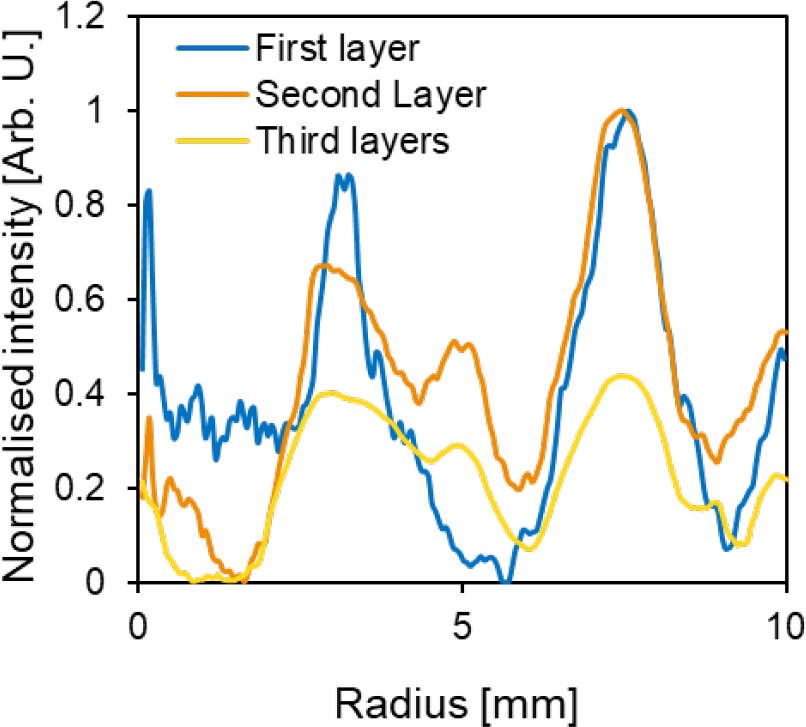
Radial profile analysis of a multilayered sample reported in Fig. 3-B.

**Figure S4.**
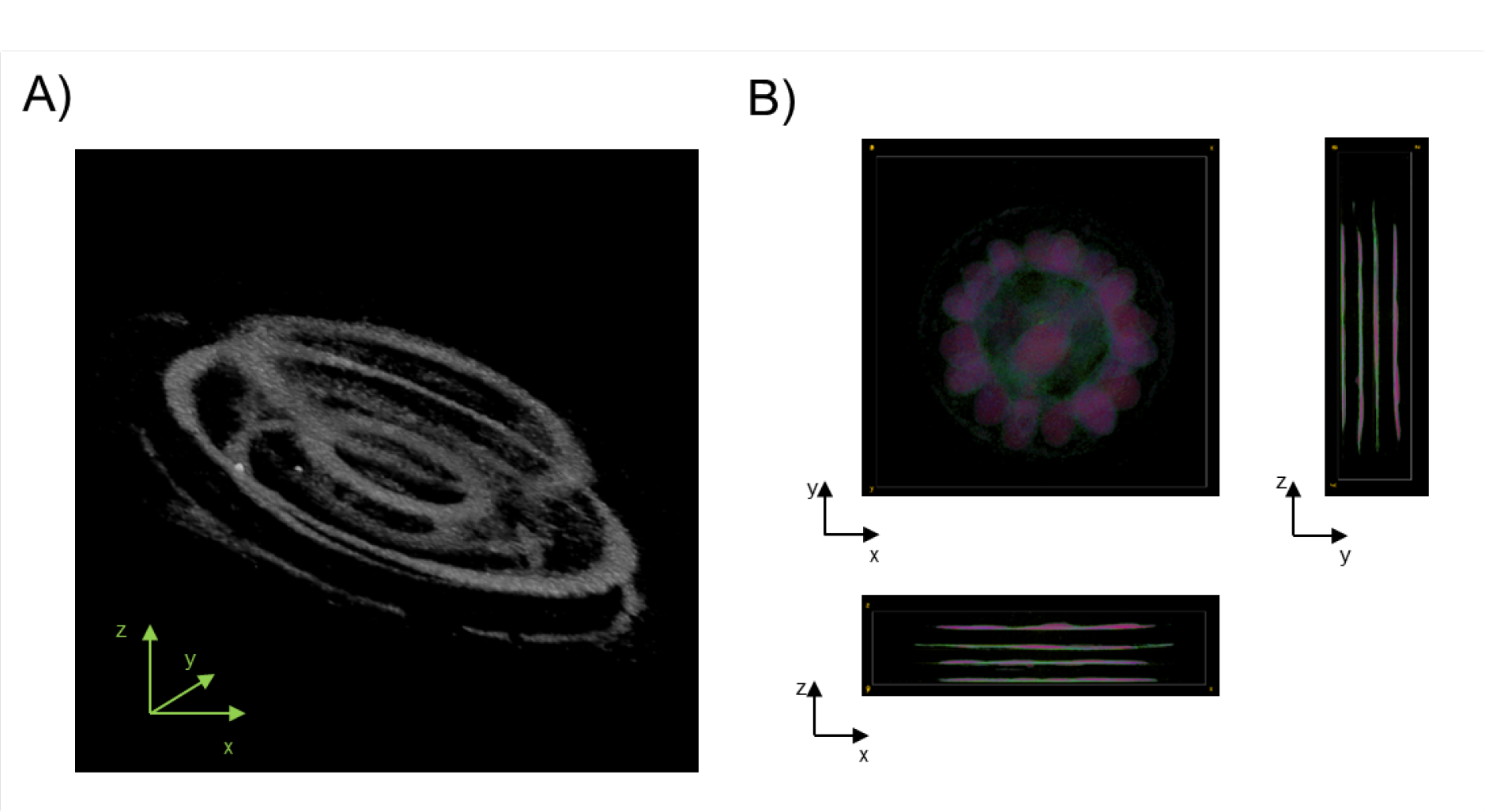
3D reconstruction (A) and 3D projection (B) of the CT scan for the multilayered sample in the circular frame.

**Figure S5.**
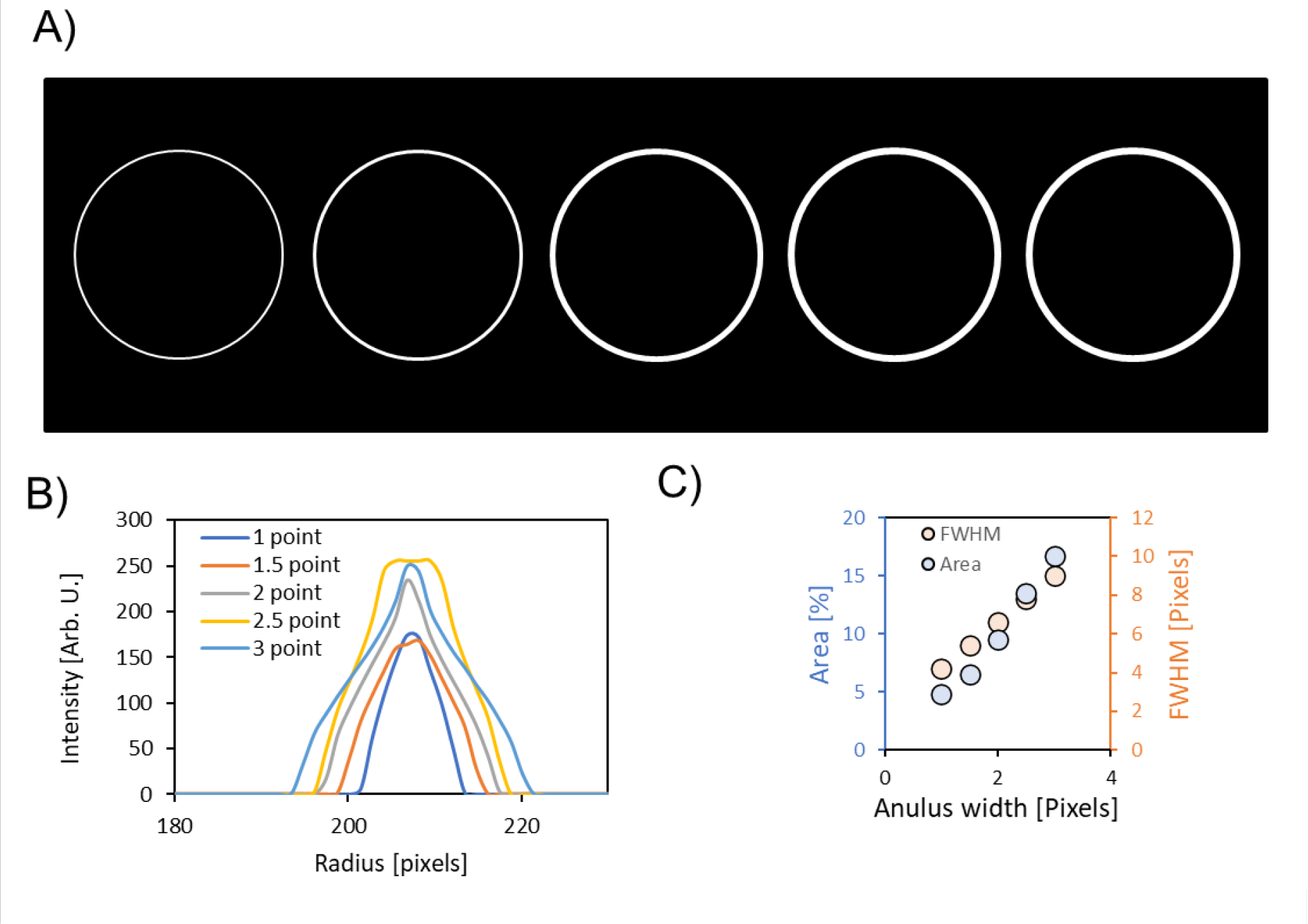
Simulated data for cell sprouting from the original annular region (A), radial profile (B) and quatification of the full width half maximum and area coverage (C).

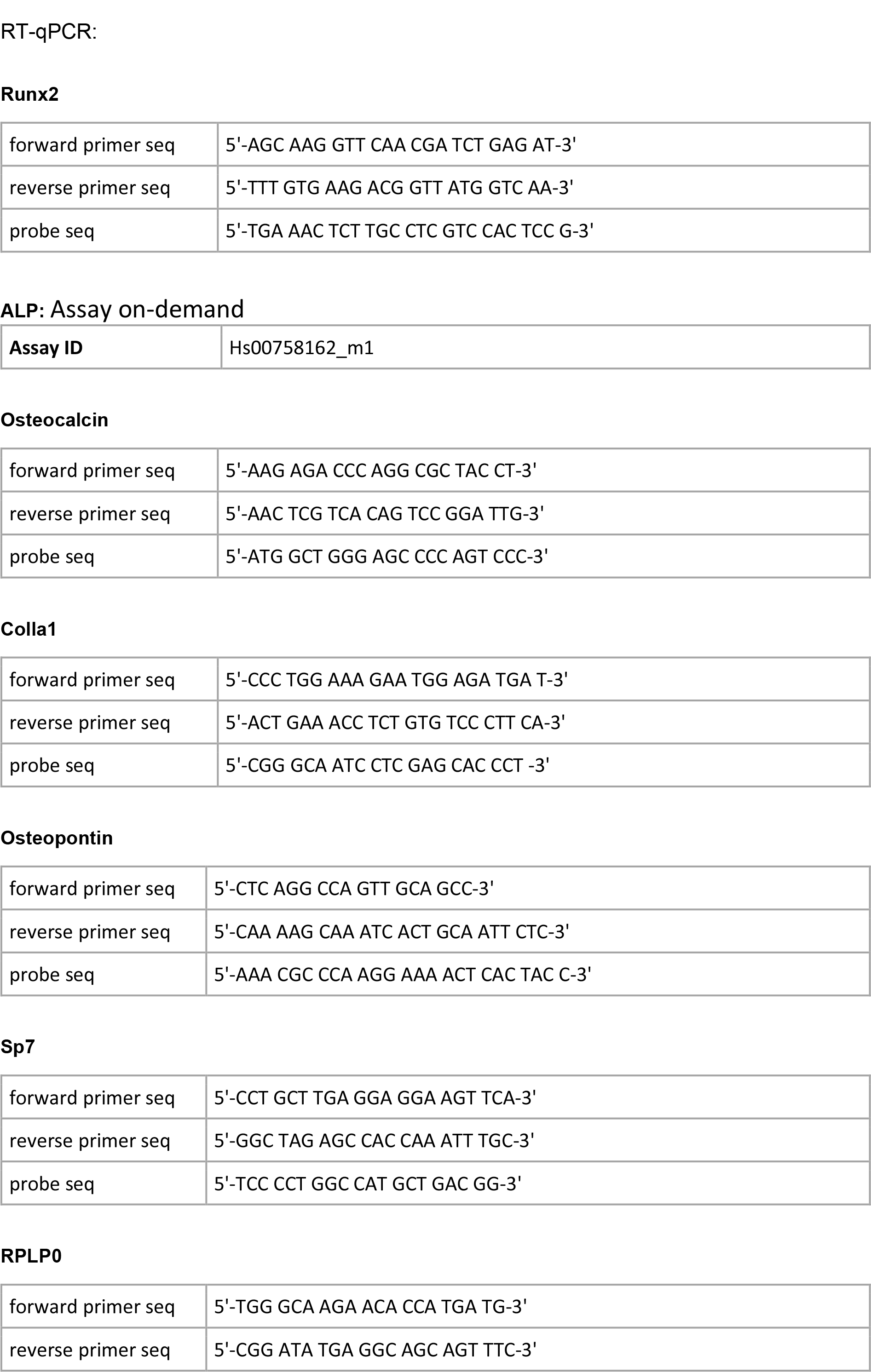
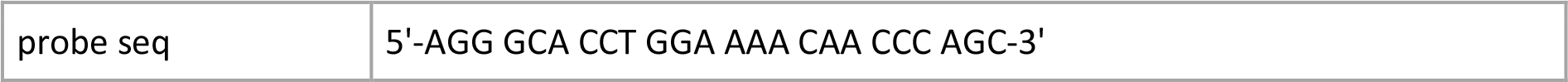

